# Clonal reproduction as a driver of liana proliferation following large-scale disturbances in temperate forests

**DOI:** 10.1101/2024.08.21.609080

**Authors:** Hideki Mori, Takashi Kamijo

## Abstract

Large-scale disturbances significantly impact forest dynamics, structure, and biodiversity. Lianas (woody vines) proliferate rapidly after such events, likely due to clonal reproduction. Understanding this process is challenging due to the need for precise disturbance history and accurate estimation of whether individuals originate from clonal reproduction, seed reproduction, or pre-existing vegetation.

This study examines whether clonal reproduction drives liana proliferation following large natural disturbances. We analyzed the dominant liana species (*Trachelospermum asiaticum* var. *asiaticum*; Apocynaceae) in temperate forests on a volcanic island. The study included young forests recovering from a volcanic eruption 22 years ago and old-growth forests unaffected by eruptions for over 800 years. We established six 10 m × 10 m quadrats (three in each forest type), divided into 1 m − 1 m grids, and sampled 587 individuals. Genetic structure was assessed using 11 newly developed nuclear microsatellite markers.

Significant clonal expansion was observed in both forest types, but stem density and genetic diversity varied markedly. Old-growth forests had 14 times more stem density and five times more genets (clones) than young forests, with greater genet intermingling and higher clonal diversity. This indicates that clonal reproduction results in high abundance and complex spatial genetic structures of the liana species in old-growth forests.

Our analysis revealed that a few genets, newly recruited via seed dispersal in early succession stages, rapidly expanded through extensive clonal reproduction, leading to long-term liana proliferation. This study highlights the importance of understanding and quantifying clonality in predicting liana population dynamics after large-scale disturbances.

## INTRODUCTION

Large-scale disturbances significantly alter the structure, dynamics, and functions of forests (Oliver 1980; Seidl et al., 2014). With climate change and forest fragmentation, these disturbances are increasing globally (Laurance, 2004; Siedl et al., 2017). Evaluating the impact of anticipated long-term increases in large-scale disturbances on forests has become a major focus in forest research (Schurman et al., 2018; McDowell et al., 2020; Patacca et al., 2023). For instance, large-scale wind disturbances and subsequent pest outbreaks have been shown to significantly restructure forest landscapes, altering canopy composition and facilitating species shifts (Schurman et al., 2018).

Lianas are crucial components of forest ecosystems and proliferate rapidly following such disturbances (Ladwig and Meiners, 2015). Events like typhoons, wildfires, and forest fragmentation can trigger long-term increases in liana abundance worldwide (Allen, 2002; Leicht-Young, 2013; Ngute et al., 2024). However, the biological mechanisms behind liana proliferation post-disturbance remain poorly understood. Lianas’ ability to quickly colonize and dominate disturbed areas often increases competition with tree species, potentially altering forest regeneration and ecosystem dynamics (Schnitzer and Bongers, 2002).

Lianas establish and colonize new environments after disturbances through seed dispersal, advanced vegetation on the forest floor (including existing seeds and seedlings), and clonal reproduction (Schnitzer and Bongers, 2002). Previous studies indicated that clonal reproduction is a major driving factor for their long-term proliferation (Yorke et al., 2013; Schnitzer et al., 2021). For instance, the increase in lianas over a decade following small-scale natural disturbances has been largely attributed to stem resprouting (Schnitzer et al., 2021). Clonal reproduction in lianas has been documented in several studies (Putz, 1984; Sakai et al., 2002; Schnitzer et al., 2012; Mori et al., 2021), highlighting its significant role in their abundance and spatial distribution patterns in forests.

The extensive clonal reproduction of plants by stolons or rhizomes not only increases their abundance but also alters their genetic diversity, including genetic structure and clonal diversity, depending on the growth strategies employed (Schmid and Harper, 1985). The “guerilla” type establishes low-density ramets over a large area, which subsequently leads to high density through intermingling of clones. Conversely, the “phalanx” type establishes and maintains high-density ramets in a small area, exhibiting a lack of intermingling and strong exclusion of other plants. Additionally, intermingling of clones has been demonstrated to enhance overall fitness in clonal plants (Matsuo et al., 2014). Therefore, an understanding of these growth types and the degree of intermingling among clones is essential for comprehending the demography and survival strategies of clonal plants, including lianas.

Despite the importance of clonal reproduction on liana proliferation, its mechanisms following large-scale disturbances are not fully understood. This is likely due to the need for precise disturbance history and the challenge of evaluating clonal reproduction alone, as established individuals may originate from clonal reproduction, seed reproduction, or pre-existing vegetation. Most existing knowledge focuses on disturbances like forest fragmentation, typhoons, and wildfires (Allen et al., 1997; Leicht-Young, 2013; Barry et al., 2015; Ngute et al., 2024), where regenerative plant propagules (e.g., soil seed banks, advanced vegetation) might not be completely eradicated. Consequently, even with a clear disturbance history, distinguishing between increases in abundance from individuals that survived the disturbance and those that newly recruited after disturbance is challenging. In contrast, volcanic eruptions, particularly those characterized by thick ash deposits, can eliminate regenerative plant propagules (Antos & Zobel, 1986; Tsuyuzaki, 1987). In such instances, the increase in abundance would be driven by propagules (e.g., seeds) introduced from external sources, making it possible to accurately estimate the origins of individual plants. Further research is necessary to unravel these complex interactions and develop a comprehensive understanding of the biological mechanisms utilized by lianas following large-scale disturbances.

This study aims to investigate whether the proliferation of liana species after large-scale natural disturbances is driven by clonal reproduction. To achieve this objective, we conducted a comparative analysis of liana abundance and genetic structure in temperate forests on a volcanic island. The volcanic island was subject to a significant volcanic eruption in 2000, which resulted in the complete loss of a substantial portion of the island’s vegetation (Fig. 1; Kamijo et al., 2008; Kawagoe et al., 2011). The study sites included young forests (shrublands) that were established after the complete loss of vegetation caused by the 2000 eruption and old-growth forests (estimated to be over 800 years old) unaffected by eruptions. A preliminary survey of vegetation changes in these two forest types indicated that lianas have recently colonized the young forests and remain consistently dominant in the old-growth forests, with spatially continuous high-density ramets on the forest floor. These distinct forest types are ideal for accurately estimating clonal reproduction in lianas by excluding the influence of pre-existing plant propagules. Specifically, we tested the following two hypotheses:

1. Does clonal reproduction primarily drive the initial colonization of liana species in young forests formed after a volcanic eruption? We hypothesize that the majority of liana individuals in these young forests are derived from clonal reproduction rather than seed reproduction due to the absence of regenerative plant propagules following the volcanic eruption.
2. How does clonal reproduction contribute to the long-term increase in abundance and genetic complexity of liana species in old-growth forests? We hypothesize that in old-growth forests, established individuals continue clonal growth, resulting in higher genetic diversity (including genetic structure and clonal diversity) and stem density compared to young forests, due to the intermingling of clones within the forest stand.

**Fig. 1.**
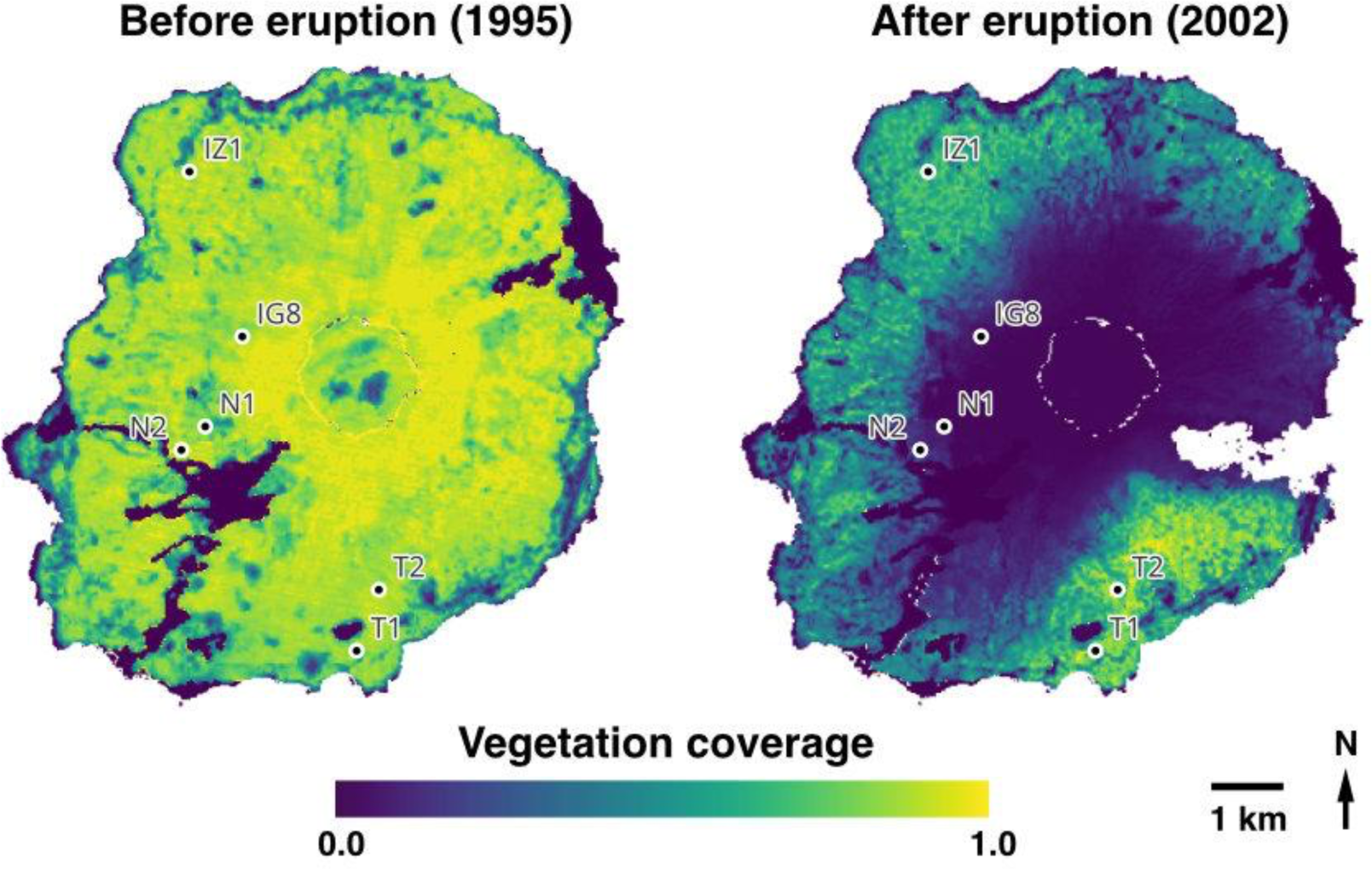
Study site locations and vegetation coverage on Miyake-jima Island before and after the 2000 volcanic eruption. The maps were created based on Yamanishi et al. (2003).

By addressing these questions, this study aims to understand the relative contributions of clonal reproduction versus seed reproduction to the rapid colonization and long-term proliferation of lianas following large-scale disturbances.

## MATERIALS AND METHODS

### Study Sites

This study was conducted on Miyake-jima Island, Tokyo prefecture, Japan (Fig. 1). Miyake-jima Island experienced a significant volcanic eruption in 2000, resulting in substantial ecological transformations (Yamanishi et al., 2003; Takahashi et al., 2008; Katoh et al., 2020; Peng et al., 2021). The volcanic eruption resulted in the complete loss of vegetation and bare land in large parts of the island (Kamijo and Hashiba, 2003; Kamijo et al., 2008; Kawagoe et al., 2011; Fig. 1). Previous research on primary succession and vegetation changes in Miyake-jima Island reported that the colonization of *Alnus sieboldiana* (Betulaceae) begins after 16 years of primary succession, forming an *Alnus* shrubland after 37 years of primary succession, and the colonization of *Castanopsis sieboldii* (Fagaceae) occurs after more than 800 years of primary succession (Kamijo et al., 2002; Table S1). The present study utilized long-term vegetation monitoring sites (Table S2; Kamijo et al., 2002; Peng et al., 2021) to select two distinct forest communities for analyzing the clonal structures of lianas, which have developed with and without large natural disturbances. The first forest type is an evergreen broadleaf forest dominated by *C. sieboldii,* which has remained unaffected by volcanic activity for over 800 years, hereafter referred to as the “old-growth forest.” The second forest type is a deciduous shrubland dominated by *A. sieboldiana,* currently in the recovery phase following the destruction of vegetation due to the 2000 eruption, hereafter referred to as the “young forest.”

### Study Species

The study species is *Trachelospermum asiaticum* (Siebold et Zucc.) Nakai var. *asiaticum* (Apocynaceae), with its nomenclature based on Yonekura and Kajita (2003). *Trachelospermum asiaticum* var. *asiaticum* is a woody climbing plant native to warm temperate forests, classified as a wind dispersed, evergreen broadleaf species. The climbing type of this species is classified as a root climber (Kusakabe et al., 2023), utilizing adhesive roots to attach to and ascend host trees, though it occasionally exhibits twining behavior around supports. On Miyake-jima Island, *T. asiaticum* var. *asiaticum* is a dominant species in the forest understory and is found across a range of forest types, from young forests to old-growth forests (Table S1; Fig. S1). In old-growth forests, it is particularly abundant, forming dense carpet-like vegetation. This species spreads across the forest floor, creating apparent individuals (ramets) while forming genetically identical clones (genets). Previous vegetation surveys in young and old-growth forests indicated that this species had established and colonized after the large natural disturbance (Table S2). Due to its remarkable clonal growth characteristics in a range of forest types, *T. asiaticum* var. *asiaticum* is an ideal species for achieving the objectives of this study.

### Sampling and DNA Extraction

Sampling was conducted at long-term monitoring sites located within two forest types, young and old-growth forests, on Miyake-jima Island in 2022 (Fig. 1). In each forest type, three quadrats (10 m × 10 m) were selected for sampling (young forest sites: N1, N2, IG8; old-growth forest sites: T1, T2, IZ1; Fig. 1), resulting in a total of six 100 m^2^ quadrats. Each quadrat was divided into 1 m x 1 m grids (totaling 100 grids per quadrat), and a ramet was randomly sampled within each grid. Lianas are known to exhibit two distinct stages of clonal growth: the on-floor stage, characterized by small individuals before climbing trees, and the on-tree stage, where individuals ascend trees (Mori et al., 2021). Therefore, sampling was conducted for both life stages, resulting in a potential maximum of 200 samples per quadrat. Following collection, samples were placed in paper bags with silica gel to maintain dryness and stored at room temperature until DNA extraction. DNA was extracted using the 2×CTAB (cetyltrimethylammonium bromide) protocol (Murray and Thompson, 1980).

### DNA Marker Development and Genotyping

Polymerase chain reaction was performed using 11 microsatellite markers designed for the study species (Table S3; Supplementary Methods S1). Sequences of *T. asiaticum* var. *asiaticum* were downloaded from the Sequence Read Archive (accession number SRR6425626) at NCBI database (http://www.ncbi.nlm.nih.gov/sra), and 11 loci of microsatellite markers (Table S3) were developed using the CMIB (CD-HIT-EST, MISA, ipcress and BlastCLUST) pipeline (Ueno et al., 2012), following the methodology described by Mori et al. (2016) (for details of marker development, see Supplementary Methods S1). The PCR products were analyzed using a 3130 Genetic Analyzer (Applied Biosystems, CA, USA). Electropherograms from each marker were meticulously examined for peak patterns using GeneMarker v1.95 (https://softgenetics.com/products/genemarker/).

### Data Analysis

Clones were identified based on the methods proposed by Arnaud-Haond et al. (2007). Firstly, the ability of the microsatellite markers to distinguish multilocus genotypes (MLGs) was tested by calculating the number of MLGs for all combinations of a given locus, and then the results were verified based on the plateaus of the genotype accumulation curves (Fig. S2). To ascertain whether stems of the same MLG belonged to the same clone, the probability of a given MLG occurring in a population under Hardy–Weinberg equilibrium (*P_gen_*) was calculated using the equation 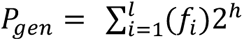 (Parks and Werth 1993), where *f_i_* is the frequency of each allele at the *i*-th locus estimated with a round-robin method, and *h* is the number of heterozygous loci. Then, the probability of obtaining *n* repeated MLGs from a population more than once by chance in *N* samples (*P*_sex_) was calculated using the equation 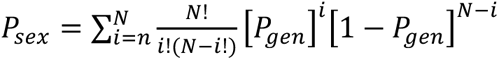 (Parks and Werth 1993). To distinguish each distinct MLG that belonged to a distinct clone, multilocus lineages (MLLs) were defined based on pairwise genetic distances. This procedure was necessary to prevent the false detection of clones due to slightly different MLGs resulting from somatic mutation or genotyping errors. The pairwise genetic distance threshold was determined with the cutoff_predictor function of R package “poppr” (Kamvar et al., 2014, 2015). MLLs are equivalent to genets (clones) in ecological studies, and we will use “genets” hereafter for MLLs. Finally, to evaluate the clonal diversity the following indices were obtained: clonal richness (*R*), Simpson’s evenness index (*V*), and the Pareto index (*β*) (Arnaud-Haond et al., 2007). Clonal richness (genotypic richness) was calculated as *R* = (*G* − 1)/(*N* − 1) where *G* is the number of genets in *N* samples. Simpson’s evenness index was calculated as *V* = (*D* – *D_min_*)/(*D_max_* – *D_min_*) where *D* is the Simpson index 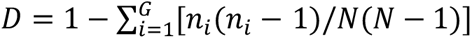, *D_min_* = {[(2*N* − *G*)(*G* − 1)]/*N*^2^} × [*N*/(*N* − 1)], *D_max_* = [(*G* − 1)/*G*] × [*N*/ (*N* − 1)], *G* is the number of unique MLGs, and *N* is the number of samples. The Pareto index is the negative value of the slope of the power law (Pareto) distribution of clonal membership. The equation for Pareto index is *N*_≥_*_x_* = *⍺X*^-*β*^ where *N*_≥_*_x_* is the number of genets containing *X* or more ramets. The Pareto index (*β*) is higher when genets have a higher density and the distribution of ramets per genet has higher evenness. Calculation of *P_gen_*, *P_sex_*, and the clonal diversity indices were conducted using the R package “RClone” ver. 1.0.3 (Bailleul et al., 2016).

Differences in liana abundance and the distribution of genets between the study sites were examined in terms of stem density, genet size, and the number of overlaps between genets. Stem density in the six quadrats was estimated using ten 1 m x 1 m sub-quadrats randomly established within a 10 m × 10 m quadrat, resulting in a total of 60 sub-quadrats in the six 100 m^2^ quadrats. The total number of stems (ramets) within ten 1 m^2^ sub-quadrats were counted for each 100 m^2^ quadrat. The size of each genet was defined as the minimum convex polygon (convex hull) enclosing the genet. The calculation of the genet size was performed using QGIS v.3.30 (QGIS Development Team, 2024). Differences of stem density and genet size among the six quadrats were tested with Tukey’s multiple comparison test using the R package “emmeans” (Russell, 2024). To test whether the total number of overlaps among genets differed among the six quadrats, the jack-knife procedure described in Mori et al. (2018) was used, with the mean and 95% confidence intervals for each quadrat being obtained from datasets generated by excluding one genet. This was done for all combinations of genets. If the contribution of a single genet to the total number of genet overlaps was significant, then a 95% confidence interval would be a high value compared to that of the mean value. We considered the values between the quadrats to be significantly different when their 95% confidence intervals did not overlap. The above-mentioned data analysis was done using R version 4.3.2 (R Core Team, 2023).

## RESULTS

### Identification of Genets

Out of 592 samples, the genotyping of 11 microsatellite loci was successfully achieved in 587 samples (99.2%) (Table 2). The genotype accumulation curves indicated that the microsatellite loci from one to five could identify all multilocus genotypes present in each quadrat (Fig. S2), demonstrating that the newly developed 11 microsatellite markers had sufficient power to identify genets (clones) for this study. The values of *P_gen_* and *P_sex_* were low (< 0.0001), suggesting that the probability of obtaining identical clones by chance was very low. Therefore, the identification of genets and the estimation of clonal structures in this study can be considered reliable.

### Clonal Structure in Young Forests

In the young forest sites (N1, N2, IG8), a total of 231 ramets were identified, with an average of 77.0 ± 28.2 per quadrat (Table 1). These ramets comprised 12 genets, with an average of 4.0 ± 1.7 per quadrat (Fig. 2; Fig. 3). The percentage of clonal ramets (those constituting multiple-ramet genets when a ramet was excluded from each genet) to the total number of ramets was 98.9 ± 1.1% on average (Table 2). These results indicate that the majority of ramets are derived from clonal reproduction, with few ramets originating from seed reproduction in the three young forest sites. At the on-floor stage, at least one ramet was found in an average of 60.7 ± 27.52 grids per quadrat, while at the on-tree stage, ramets were found in an average of 16.3 ± 2.5 grids per quadrat (Table 1), indicating that lianas were approximately four times more frequently found at the on-floor stage than the on-tree stage. The average number of ramets per 1 m^2^ was 10.5 ± 10.4, with no significant differences among the young forest sites (Fig. 3). The average size of genets was 28.9 ± 32.0 m^2^ (Fig. 3; Fig. S5), with no significant differences among the three young forest sites (Fig. 3). There were also no significant differences in the number of overlaps between genets among the three young forest sites (Fig. 4). These results indicate that the patterns of stem density, genet size, and genet overlaps are similar and consistent across the young forest sites.

**Fig. 2.**
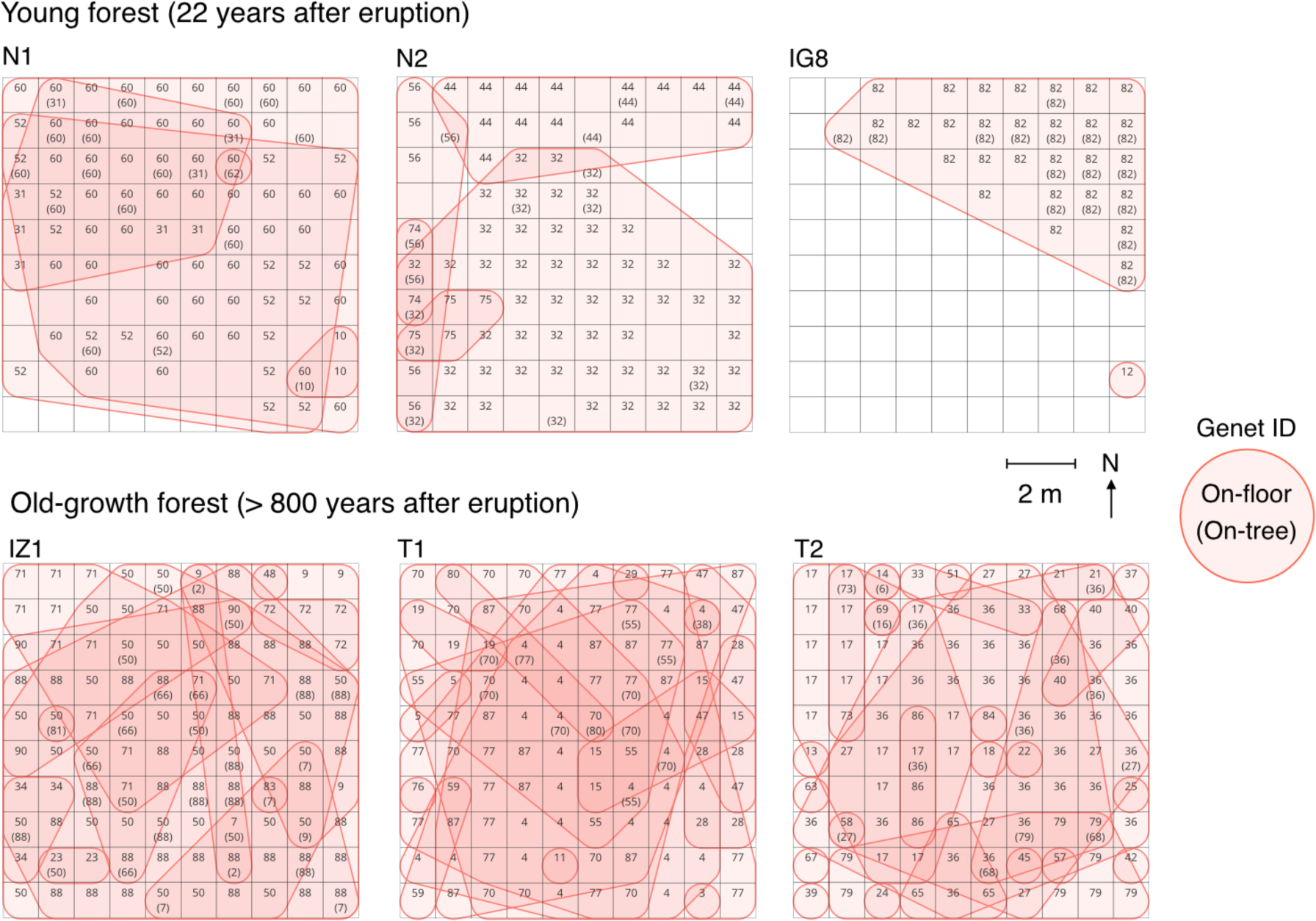
Distribution map of ramets and genets in the six 10 m × 10 m quadrats. Each quadrat is divided into 1 m grids (N = 100). Numbers in the grids represent unique identifiers for genets. Numbers are shown with parentheses when ramets were at the on-tree stage. The shapes of genets are shown with solid polygons.

**Fig. 3.**
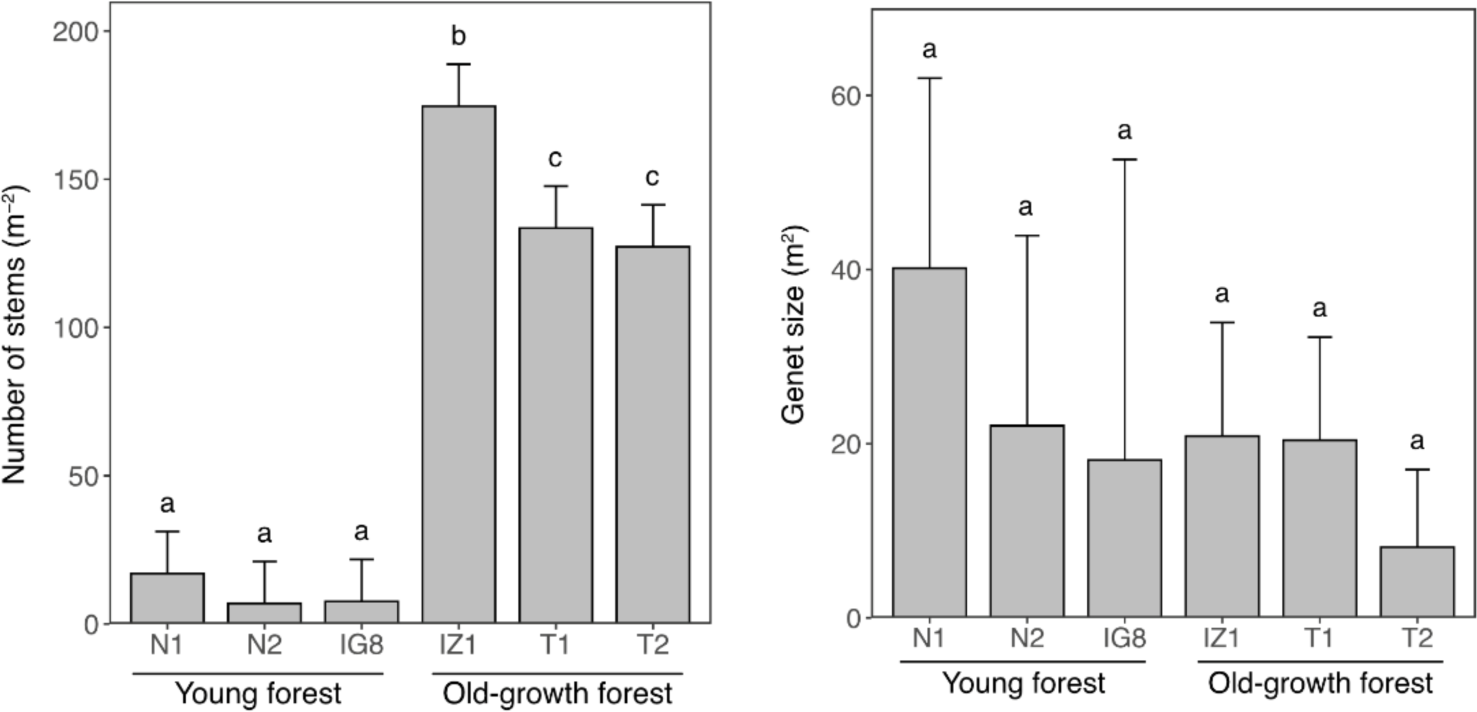
The number of stems (left panel) and the size of genets (right panel) in the study sites. Mean values with the corresponding 95% confidence interval are shown for the six quadrats. Different letters indicate significant differences based on the Tukey comparison test (confidence level = 0.95).

**Fig. 4.**
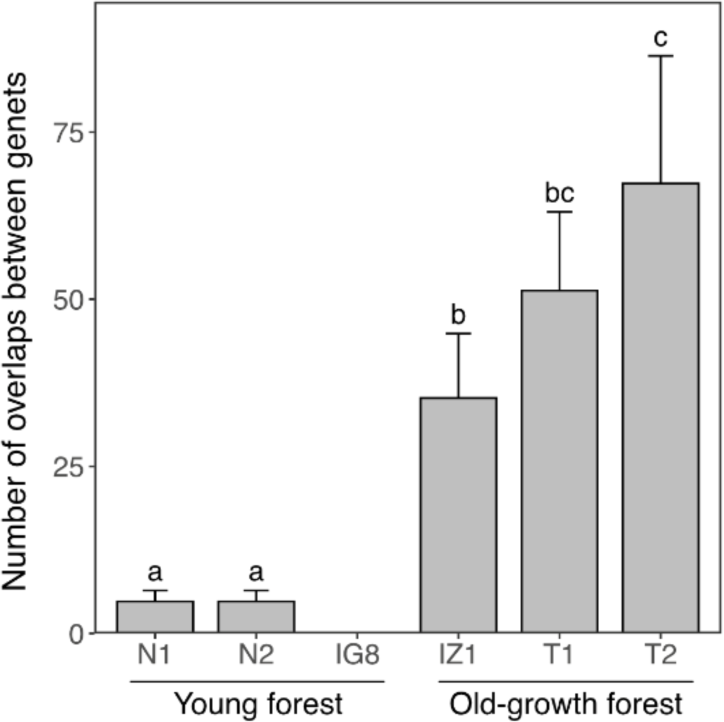
The number of overlaps between genets in each study site. Variability of the genet overlaps in the study sites was calculated by generating datasets that excluded one genet from all combinations. Error bars represent a 95 % confidence interval for the generated datasets. Different letters indicate that the 95% confidence intervals did not overlap between the study sites.

**Table 1.**
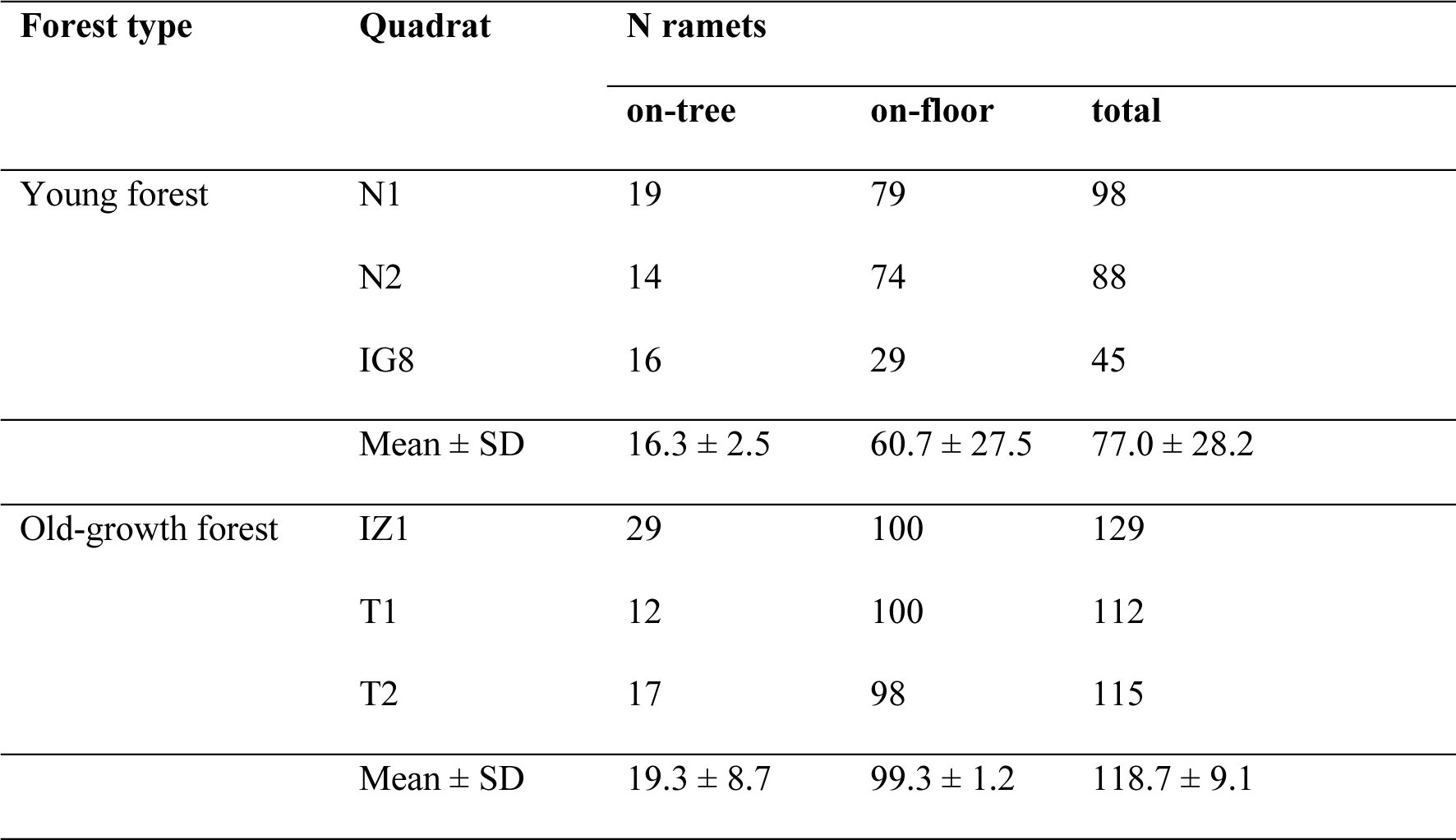
The number of ramets sampled and genotyped in this study. The ramets which climbed tree (“on-tree” column) and the ramets which had not yet started climbing (“on-floor” column) were randomly sampled, one per 1 m × 1 m grid (N = 100) of six quadrats.

### Clonal Structure in Old-Growth Forests

In the old-growth forest sites (IZ1, T1, T2), a total of 356 ramets were identified, with an average of 118.7 ± 9.1 per quadrat (Table 1). These ramets comprised 61 genets, with an average of 20.3 ± 8.5 per quadrat (Fig. 2; Fig. 3). The percentage of clonal ramets to the total number of ramets was 91.3 ± 8.6% on average (Table 2). Similar to the young forest sites, these results indicate that the majority of ramets are derived from the clonal reproduction in the three old-growth forest sites. At the on-floor stage, at least one ramet was found in an average of 99.3 ± 1.2 grids per quadrat, while at the on-tree stage, ramets were found in an average of 19.3 ± 8.7 grids per quadrat (Table 1). This indicated that lianas were approximately five times more frequently found at the on-floor stage than the on-tree stage. The average number of ramets per 1 m^2^ was 145.1 ± 36.1, with significant differences found between IZ1 and the other two sites (Fig. 3). The average size of genets was 14.5 ± 23.2 m^2^ (Fig. 3; Fig. S5), with no significant differences among the three sites (Fig. 3). There was a significant difference in the number of overlaps between IZ1 and T2 (Fig. 4). These results indicate that while overall tendencies in genet size were similar, there were differences in stem density and the frequency of genet overlaps within old-growth forests.

**Table 2.**
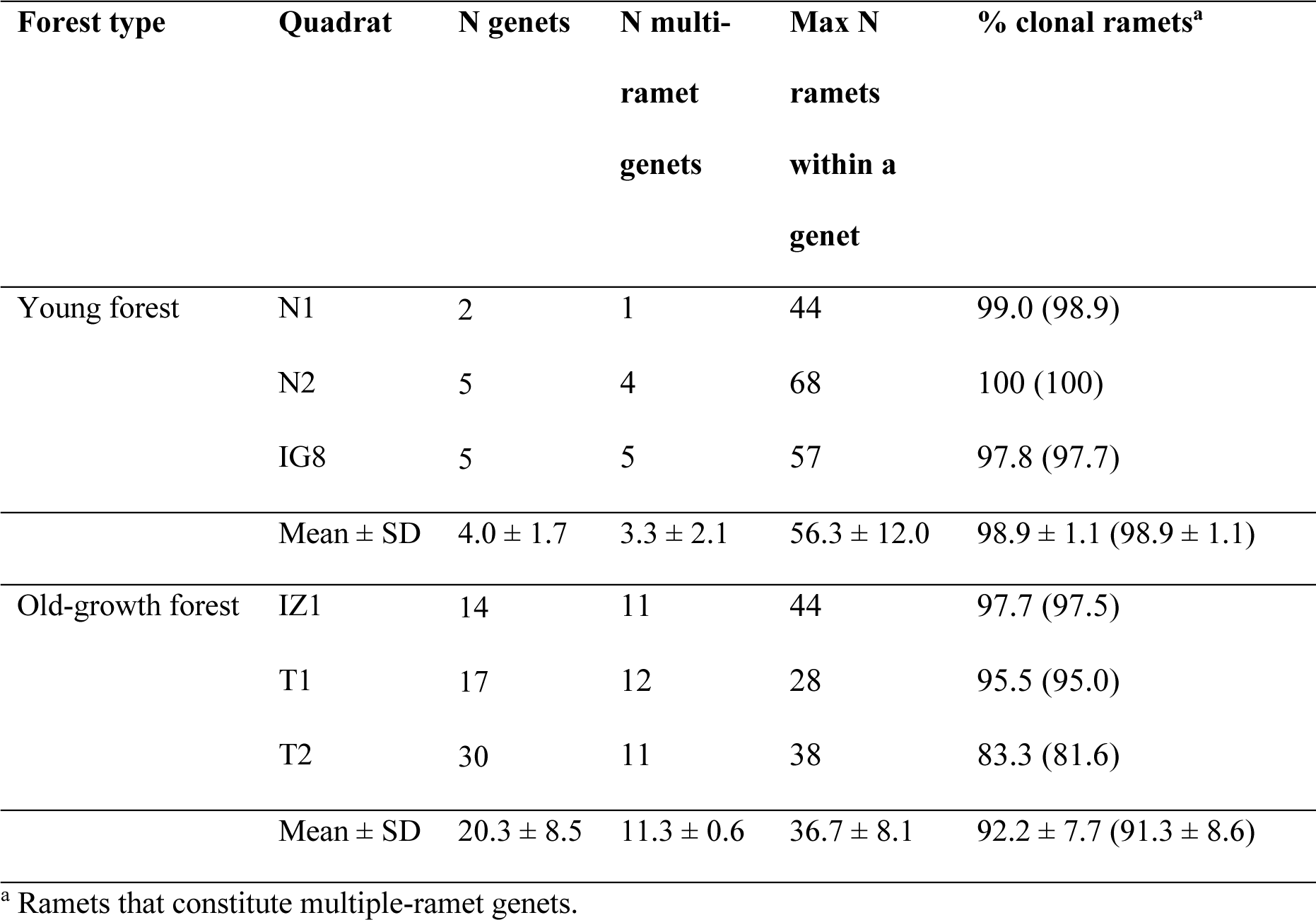
Summary of the number of genets and ramets identified in this study. Number in parentheses represents the percentage of clonal ramets when a ramet was excluded from each genet.

### Comparison of Clonality between Young and Old-Growth Forests

The clonal diversity indices (*R, V, β*) were consistently lower in the young forest sites compared to the old-growth forest sites (Table 3). This suggests that the clonality of the study species in the old-growth forests is characterized by higher richness and greater evenness. The number of ramets (stem density) was significantly higher in all old-growth forest sites than in the young forest sites (Fig. 3; Wilcoxon rank-sum test, P < 0.001; Fig. S5). Specifically, the stem density in the old-growth forests was over 14 times higher than in the young forests, indicating that the clonal structure in old-growth forests is characterized not only by higher clonal diversity but also by greater abundance compared to young forests.

**Table 3.**
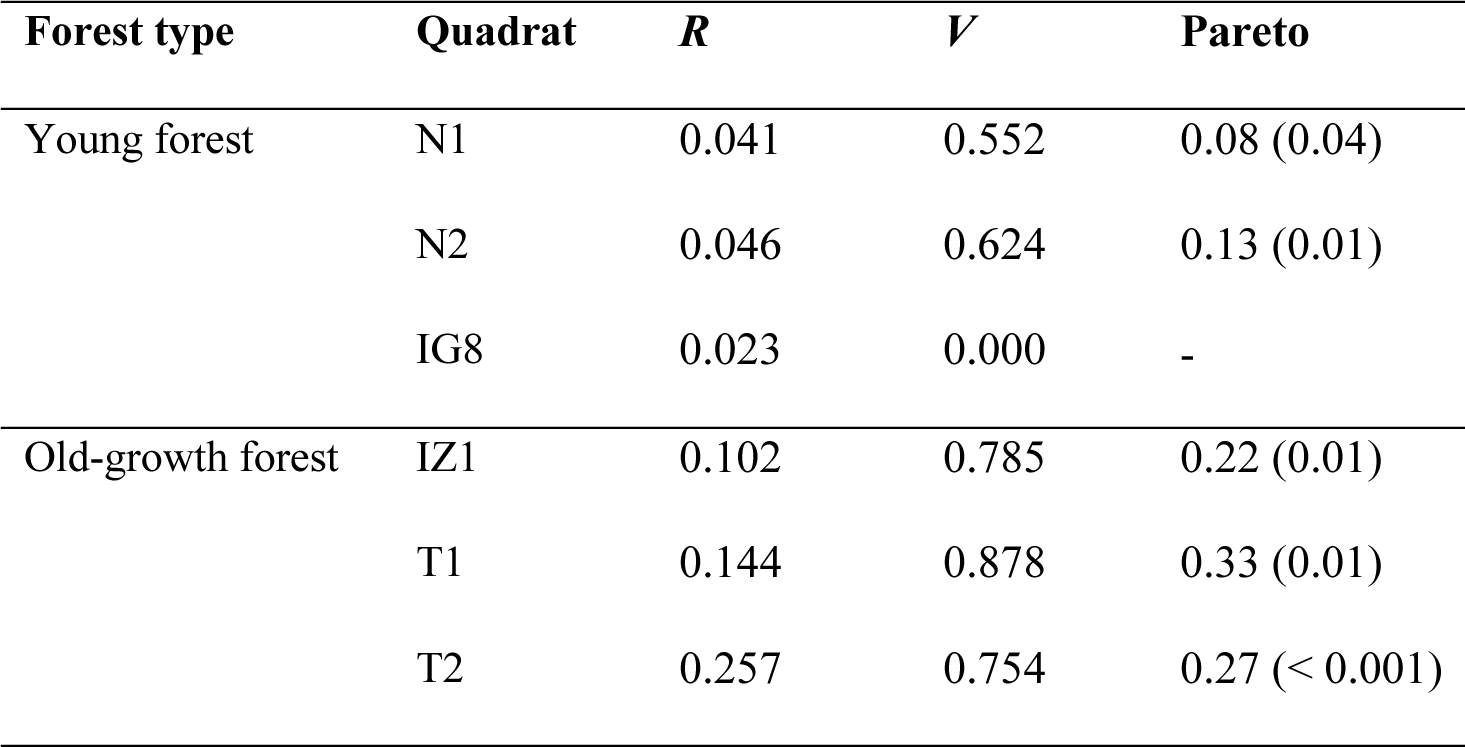
Summary of clonal diversity indices: clonal richness (*R*), Simpson’s evenness index (*V*), and Pareto index. Number in parentheses represents P values. Pareto index for IG8 was not calculated due to the very few number of genets (N=2).

| Forest type | Quadrat | $R$ | $V$ | Pareto |
| --- | --- | --- | --- | --- |
| Young forest | N1 | 0.041 | 0.552 | 0.08 (0.04) |
|  | N2 | 0.046 | 0.624 | 0.13 (0.01) |
|  | IG8 | 0.023 | 0.000 | - |
| Old-growth forest | IZ1 | 0.102 | 0.785 | 0.22 (0.01) |
|  | T1 | 0.144 | 0.878 | 0.33 (0.01) |
|  | T2 | 0.257 | 0.754 | 0.27 (< 0.001) |

There were no significant differences in the size of the genets between the two forest types (Fig. 3; Wilcoxon rank-sum test, P = 0.46; Fig. S5). However, the number of overlapping genets were generally higher in the old-growth forests compared to the young forests (Fig. 4). These results indicate that, while the genet size due to extensive clonal reproduction was similar between the sites, the clonal structure in the old-growth forests was characterized by higher overlaps of genets compared to the young forests. The higher overlaps of genets in the old-growth forests were further supported by the fraction of pairs of ramets sharing the same genet, which was consistently higher in young forests (0.66 ± 0.23) compared to old-growth forests (0.18 ± 0.07) up to a spatial distance of seven meters (Fig. S6).

## DISCUSSION

This study evaluated the clonal proliferation processes of the liana species *T. asiaticum* var*. asiaticum* following a large natural disturbance by comparing populations in young forests established soon after a volcanic eruption with old-growth forests unaffected by the eruption. The findings suggest that clonal reproduction substantially contributed to the rapid colonization of the liana species after the large natural disturbance, and that clonal reproduction also played a key role in long-term increases in abundance, clonal diversity and complexity in genetic structure.

### Rapid Colonization with Clonal Proliferation after Large Natural Disturbance

It was confirmed that nearly all individuals (ramets) were derived from clonal reproduction, averaging 98.9% in young forests. This finding suggests that the colonization of lianas in young forests is predominantly driven by clonal reproduction. Previous studies have reported that lianas rapidly colonize following large-scale natural disturbances such as typhoons and wildfires (Allen et al., 2005; Leicht-Young et al., 2013), however, these disturbances do not eradicate regenerative plant propagules. Therefore, the results from those studies reflect the combined effects of surviving liana individuals, seed reproduction (seed dispersal, seed bank), and clonal reproduction, making it challenging to quantitatively assess the impact of clonal reproduction on liana proliferation. In contrast, the young forests in this study were established after the volcanic eruption that destroyed the vegetation (Yamanishi et al., 2003; Takahashi et al., 2008). Hence, the apparent increase in individual numbers through clonal growth observed in this study reflects the colonization by new individuals (seed-originated) that dispersed from the surrounding environment and subsequently proliferated through clonal growth, rather than from pre-existing seeds and advanced vegetation before the disturbance. The significant increase in lianas due to clonal reproduction in young forests suggests that the rapid proliferation following large disturbances reported in other forest communities (e.g., Allen et al., 1997; Barry et al., 2015) could also largely be attributed to clonal reproduction. This finding underscores the importance of considering clonality when understanding liana proliferation after disturbances.

The low number of genets found in the young forest sites (an average of four genets per quadrat) suggests that the study species increased its abundance through clonal reproduction from a few seed-originated individuals. In addition, these individuals likely dispersed and established within the last ten years, based on the absence of the species in these sites ten years prior to the sampling for this study (Table S2). The extensive vegetation decline caused by the eruption necessitates seed dispersal from the surrounding environment essential for establishment in young forest sites (Antos & Zobel, 1986; Wood & del Moral, 1987; Nishi & Tsuyuzaki, 2004). However, despite this necessity, the contribution of seed reproduction to the overall abundance increase after new colonization in young forests was relatively small. Although seed establishment should be possible in young forest sites (shrublands) with abundant light resources, the relatively low number of seed-originated individuals may be due to limited seed sources in the surrounding area and/or other environmental factors such as soil nutrient limitations (Eriksson & Ehrlén 1992; López et al., 2010; Peng et al., 2021). To further clarify these factors, it is necessary to elucidate the life history strategy of seed reproduction by long-term monitoring of the survival and environmental response of lianas at seed to seedling stages (Mori et al., 2019).

### Long-Term Increase in Abundance and Genetic Complexity by Clonal Reproduction

The large number of individuals observed in the old-growth forest sites indicates that the liana species had increased its stem density through long-term clonal reproduction, forming a complex genetic structure. Specifically, stem density per square meter in the old-growth forests was 14 times higher than in the young forests, highlighting the significance of clonal reproduction in the long-term increase in liana abundance in closed canopy forests. The contribution of clonal reproduction to the increase in liana abundance was consistent with previous findings (Yorke et al., 2013; Schnitzer et al., 2021). In the tropical old-growth forests of Costa Rica, a study reported that over an eight-year period, the number and basal diameter of lianas increased by 15% to 20%, with part of this increase attributed to clonal growth (Yorke et al., 2013). Additionally, in the tropical forests of Barro Colorado Island, Panama, the number of liana stems increased by approximately 30% over ten years following small-scale natural disturbances, with many of these increases attributed to clonal coppiced stems (Schnitzer et al., 2021). This indicates that clonal reproduction plays a central role in the long-term proliferation of lianas in various forest communities.

The significantly larger abundance found in the old-growth forests compared to young forests in this study could have been due to the extensive overlapping (intermingling) of genets derived from clonal reproduction. The high level of intermingling of genets indicates that the study species exhibits the “guerilla” type growth form (Schmid and Harper, 1985). Clonal plants are known to form high-density and complex spatial genetic structures through the intermingling of genets with extensive clonal growth (e.g., Kitamura et al., 2001; Reisch et al., 2007; Ohsako, 2010). This characteristic of clonal growth could have decreased the cost of clonal reproduction and increased reproductive fitness, as reported in a dwarf bamboo (Matsuo et al., 2014). In the old-growth forest sites, the dark light environment and high ramet density on the forest floor likely made seedling establishment through seed reproduction difficult. However, compared to the young forest sites, the larger number of genets (5 times on average) were identified in the old-growth forest sites. These findings suggested that both continuous new recruitment through seed reproduction and an increase in stem density through clonal reproduction had shaped the spatial genetic structure and abundance of the liana species in the old-growth forests. Indeed, large-scale disturbances have been shown to significantly alter the abundance and genetic structure and diversity in forest plants (Jackson and Fahrig, 2014; Davies et al., 2016; Soares et al., 2019). Furthermore, prolonged clonal growth of herbaceous plant populations has been recognized as a key factor enabling long-term persistence following such disturbances, including habitat fragmentation in forests (Honnay et al., 2005). These findings indicate that recruitment through seed reproduction, combined with extensive clonal reproduction, shapes the genetic structure and diversity of liana populations, with clonal reproduction playing a more dominant role for the long-term liana proliferation following large disturbances.

## CONCLUSIONS

This study demonstrated that clonal growth is crucial for the rapid and long-term increase in abundance (stem density) and genetic diversity (genetic structure and clonal diversity) of the liana species after large-scale natural disturbances. The processes suggested by this study are as follows:

1. After the volcanic eruption, a few genets established via seed dispersal into the recovering *Alnus* shrublands increased their abundance through extensive clonal growth on the forest floor.
2. Over time, as the forests transitioned into climax communities dominated by old-growth *C. sieboldii*, clonal growth significantly increased the abundance of the liana species, resulting in high-density ramets and complex spatial genetic structures.

Thus, *T. asiaticum* var. *asiaticum* exhibits both pioneer characteristics and high shade tolerance. It rapidly establishes in bright shrublands through both seed and clonal reproduction, while achieving long-term increases in abundance through clonal growth in closed-canopy old-growth forests. These findings indicate that the combination of clonal and seed reproduction enables the liana species to adapt effectively to forest succession and environmental changes following large-scale disturbances.

Increasing liana dominance has primarily been reported in tropical forests (Ngute et al., 2024), partly due to their high diversity and abundance in these regions. However, this study suggests that clonal reproduction can significantly enhance liana proliferation after large-scale disturbances in higher latitudes, such as temperate regions. As large-scale disturbances increase worldwide (Siedl et al., 2017), lianas may continue to proliferate across various forest types. Given their structural and functional importance in forest ecosystems and significant impact on community dynamics, lianas are increasingly considered in future predictions of forest communities (van der Heijden et al., 2013; Meunier et al., 2021). This study underscores the necessity of evaluating the clonal nature of lianas to comprehensively understand and predict changes in their abundance and genetic diversity caused by forest disturbances.

## Acknowledgements

We thank Mr. S. Honma and Ms. M. Muto (University of Tsukuba) for their help during the field work. We thank Dr. S. Ueno (FFPRI) for his advice on SSR marker development. We also thank Ms. C. Furusawa and other staff members in the Department of Forest Molecular Genetics and Biotechnology, FFPRI for the molecular experiments during this study. This study was supported by JSPS KAKENHI Grant Numbers 21K14882, 24K01802. In this study, we utilized the supercomputer of AFFRIT, MAFF, Japan.

## Author Contributions

HM and TK conceived the study design and conducted the field survey. HM performed the molecular experiments, analyzed the data, and wrote the manuscript, with TK providing editorial advice.

## Data Availability Statement

The datasets used and/or analyzed during the current study are available from the corresponding author on reasonable request.

## Notes

### Competing Interest Statement

The authors have declared no competing interest.

