## Supplementary material for "Clonal reproduction as a driver of liana proliferation following large-scale disturbances in temperate forests": Fig. S1

**Fig. S1.** Photographs of the study species in the two forest types of the study sites.

Young forest

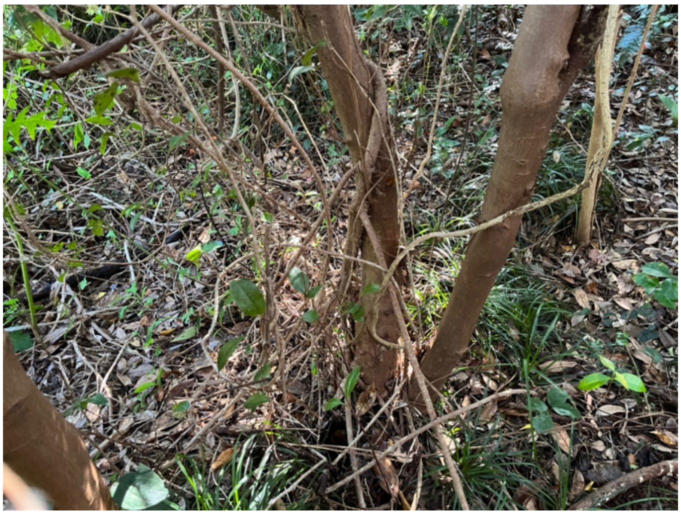

Old-growth forest

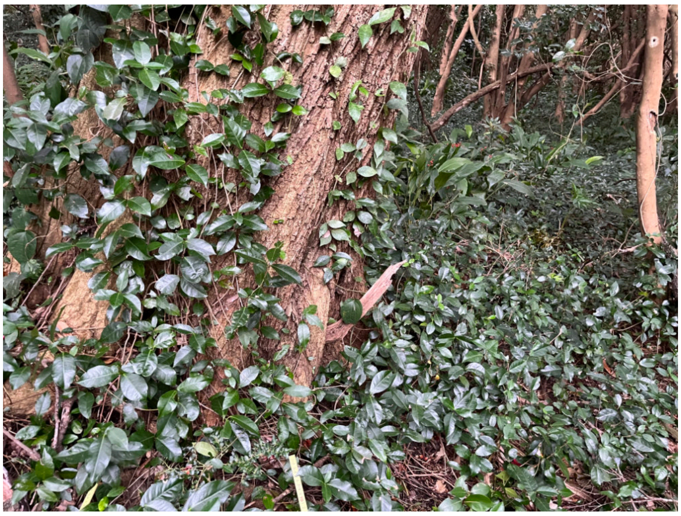
