## Supplementary material for "Clonal reproduction as a driver of liana proliferation following large-scale disturbances in temperate forests": Fig. S2

**Fig. S2.** Genotype accumulation curves of the 11 SSR markers developed in this study. Horizontal dashed line indicates the maximum number of multilocus genotypes identified in the study site (MLG).

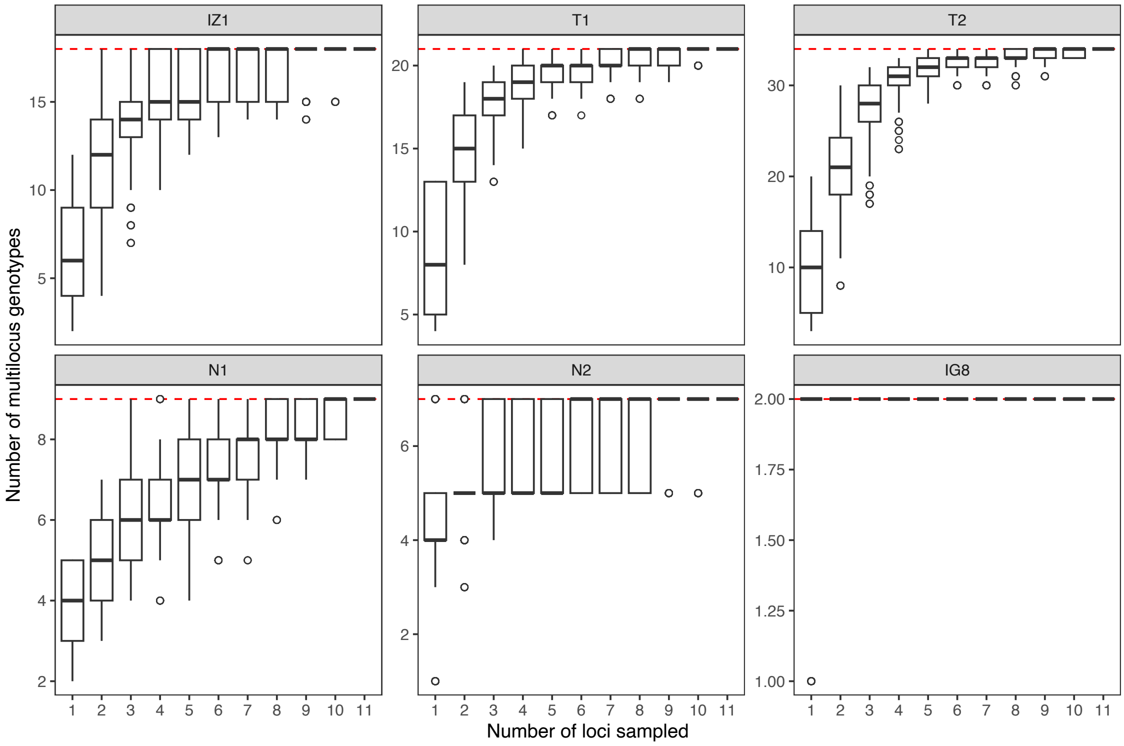
