## Supplementary material for "Clonal reproduction as a driver of liana proliferation following large-scale disturbances in temperate forests": Fig. S3

**Fig. S3.** Map of clonal structure in the six 10 m x 10 m quadrats. Each quadrat is divided into 1 m grids. Numbers and colors represent genets on the forest floor. Numbers in the parenthesis represent ramets that were on tree. Shaded grids indicate samples which were not successfully genotyped.

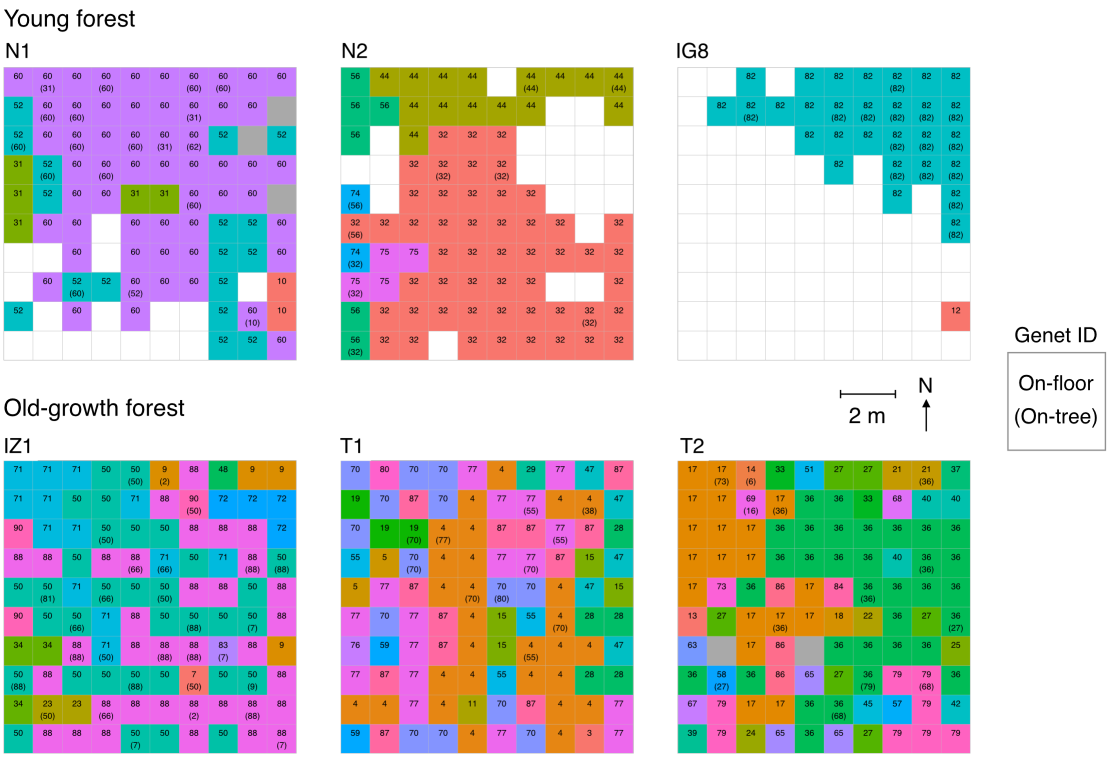
