## Supplementary material for "Clonal reproduction as a driver of liana proliferation following large-scale disturbances in temperate forests": Fig. S4

**Fig. S4.** Number of genets that were found only on the forest floor ("floor"), only on trees ("tree"), and both on the forest floor and on trees ("both").

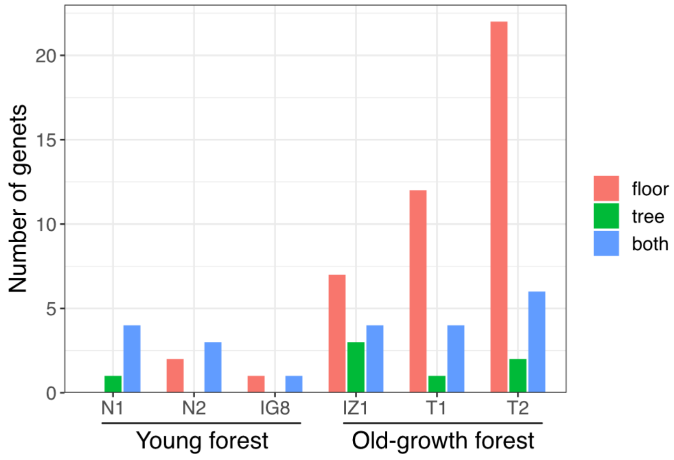
