## Supplementary figures and images for "Clonal reproduction as a driver of liana proliferation following large-scale disturbances in temperate forests"

### Fig. S5

**Fig. S5.** Genet size and stem counts in young and old-growth forest sites.

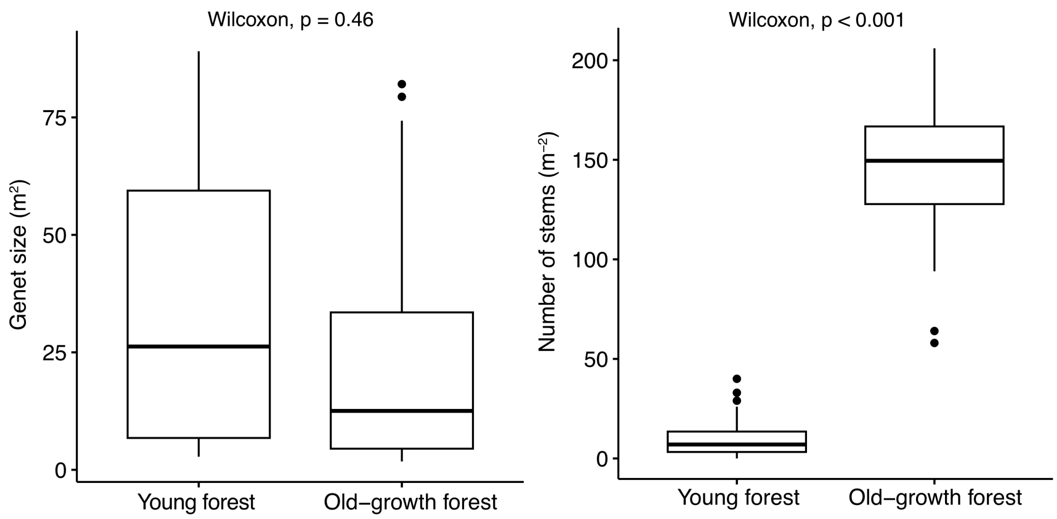
