## Supplementary material for "Clonal reproduction as a driver of liana proliferation following large-scale disturbances in temperate forests": Fig. S6

**Fig. S6.** Spatial distance and the corresponding probabilities of clonal identity (the fraction of pairs of ramets sharing the same genet; Fr) in the study sites.

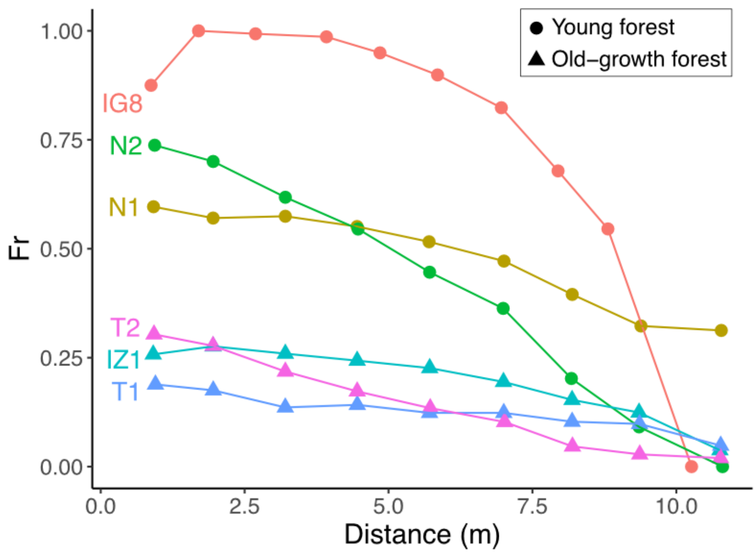
