## Supplementary material for "Clonal reproduction as a driver of liana proliferation following large-scale disturbances in temperate forests": Table S1

**Table S1.** Stages of primary succession and vegetation changes over time in Miyakejima Island. Table was created based on Kamijo et al. (2002).

| Stage (yr-old) | Primary Succession Process |
|---|---|
| 0 | Bare land |
| 16 | Colonization of *Alnus* and *Reynoutria* |
| 37 | *Alnus* shrub |
| - | Facilitation by N-fixation of *Alnus*.<br>Colonization of *Prunus* and *Machilus*.<br>Rapid above-ground-biomass accumulation. |
| 125 | *Machilus* and *Prunus* forest |
| - | Disappearance of *Alnus* and *Prunus*.<br>Colonization of *Castanopsis*. |
| > 800 | *Castanopsis* forest |
