## Supplementary material for "Clonal reproduction as a driver of liana proliferation following large-scale disturbances in temperate forests": Table S2

**Table. S2.** Change in abundance of the study species on the forest floor in the six quadrats in 2012 and 2020 based on Braun-Blanquet cover-abundance scale.

| Year | Young forest | | | Old-growth forest | | |
|---|---|---|---|---|---|---|
| | IG8 | N1 | N2 | T1 | T2 | IZ1 |
| 2012 | + | | | 44 | 44 | 44 |
| 2020 | 11 | 22 | 22 | 44 | 44 | 44 |
