## Supplementary material for "Clonal reproduction as a driver of liana proliferation following large-scale disturbances in temperate forests": Table S3

**Table. S3.** Characteristics of 11 microsatellite loci for *Trachelospermum asiaticum* var. *asiaticum.*

| **ID** | **Forward primer sequence** | **Reverse primer sequence** | **SSR motif sequence** | **PCR Product size range (bp)** | ***Na*** | ***Ne*** | ***Ho*** | ***He*** |
|---|---|---|---|---|---|---|---|---|
| Ta01 | GCCTTGCCAGCCCGCAAAGTAG GGAGAGGAGGGAGTGGC | GTTTCTTACCAACACTT CATTTCATCCAAGGC | (AG)13 | 95-145 | 6 | 3.760 | 0.500 | 0.734 |
| Ta02 | CAGGACCAGGCTACCGTGACTG AGTGGAATACAGGAGGTCTTC | GTTTCTTGCTCGTAAAC CCTGCAACCCAAC | (CT)13 | 135-170 | 4 | 1.907 | 0.448 | 0.476 |
| Ta03 | CGGAGAGCCGAGAGGTGTGGT GGGATGATGTAGTGGGTGG | GTTTCTTAAACTAAGCA CCACCAACCGCGC | (TC)18 | 160-250 | 21 | 14.885 | 0.897 | 0.933 |
| Ta04 | GCCTTGCCAGCCCGCAGTCTTG CCTCTTCAGAATCTGGTG | GTTTCTTGTCATTCCAC AAGAGAGACTGGTC | (TC)25 | 90-150 | 8 | 5.455 | 0.900 | 0.817 |
| Ta05 | GCCTTGCCAGCCCGCTTCAGGT TGTGAAGTGTGGCATC | GTTTCTTGCTCAAGTGC TCCAGGACAAGAC | (TA)14 | 140-200 | 4 | 2.985 | 0.700 | 0.665 |
| Ta06 | CAGGACCAGGCTACCGTGCTCG AGTATCATTCATTTGGCAAGG | GTTTCTTAAATCAGTTA CGACAGAGGGTGC | (TA)13 | 180-240 | 3 | 1.474 | 0.241 | 0.322 |
| Ta07 | CGGAGAGCCGAGAGGTGCCTG GATCTTTGCATTTGAACTTGCC | GTTTCTTAAATGATAGA GAGTAGCACACTGTCC | (AG)13 | 130-170 | 7 | 4.865 | 0.733 | 0.794 |
| Ta08 | GCCTCCCTCGCGCCATCCAAAT CACCAGTTCACCACACAG | GTTTCTTGCATGTTGTTA CTGTATGTTGGC | (AT)28 | 90-110 | 6 | 2.885 | 0.667 | 0.653 |
| Ta09 | GCCTTGCCAGCCCGCAACGAAT CACCGTGACTGGCAG | GTTTCTTGCATCGCTTG GCATTTGTGC | (AT)19 | 157-190 | 9 | 6.294 | 0.733 | 0.841 |
| Ta10 | GCCTTGCCAGCCCGCTCGCTCA CCTTAGCTTAGTAAATCAC | GTTTCTTCTCCATCTCTC ACTCTCGGCTAC | (GA)23 | 166-196 | 11 | 2.711 | 0.533 | 0.631 |
| Ta11 | GCCTTGCCAGCCCGCACACGTA TGATTTGACCCATGCCC | GTTTCTTCATGCATACG TAAGCTAAATTCCGG | (TA)14 | 125-150 | 4 | 2.278 | 0.800 | 0.561 |
