## Supplementary Methods S1 for "Clonal reproduction as a driver of liana proliferation following large-scale disturbances in temperate forests"

**Supplementary Methods S1: Methods for the development of 11 microsatellite markers**

Genomic sequence data for *Trachelospermum asiaticum* var. *asiaticum* were downloaded from the Sequence Read Archive (accession number SRR6425626) at NCBI (http://www.ncbi.nlm.nih.gov/sra). The total amount of sequence data was 225.1 megabase pairs (Mbp). Reads were processed with Trim Galore! version 0.4 (Krueger 2015) and were assembled with PEAR version 0.9.8 (https://cme.h-its.org/exelixis/web/software/pear/) using default parameters. All the assembled sequences were used for the CMIB (CD-HIT-EST, MISA, ipcress and BlastCLUST) pipeline (Ueno et al. 2012) to design primers with the number of repeat units ≥6, 5, 4, 3, and 3 for di-, tri-, tetra-, penta-, and hexa-simple sequence repeats (SSRs), respectively. Primer pairs were BLASTed against the NCBI nr database with an e-value of 1e-3. We selected 41 primer pairs for SSRs with the number of repeat units ≥10 and based on the BLAST hits. Forward primers with tail sequences (Blacket et al. 2012) and reverse primers with ‘pig-tail’ (Brownstein et al. 1996) were synthesized by Eurofins Genomics (Tokyo, Japan).

Leaf samples from 30 individuals along the roadside were collected in Miyake-jima island of Tokyo prefecture, Japan. All the samples were collected at least 50-m apart to avoid collecting clonally reproduced stems.

DNA was extracted from leaf tissue (10 mg) from 30 samples using a modified CTAB protocol (Tsumura et al. 1995). The initial polymerase chain reaction (PCR) was performed for two individuals in a 10-μL volume containing 1X Multi-plex PCR master mix (Qiagen), primer mix, and 5–10 ng of template DNA. The primer mix contained both forward and reverse primers and one of the fluorescently labeled TAIL primers (Blacket et al. 2012). PCR was performed using the following thermal profiles: 15 min at 95 °C, followed by 35 cycles of 30 s at 94 °C, 90 s at 60 °C, 60 s at 72 °C, and then a final extension step at 60 °C for 30 min. The products were analyzed using a 3130 Genetic Analyzer (Applied Biosystems, CA, USA) with GeneScan 600 LIZ size standard (Thermofisher, MA, USA). Electropherograms from each marker were critically checked for clear peak pattern using Fragman version 1.0.9 (Covarrubias-Pazaran et al. 2016). PCR amplification and genotyping for the rest of the samples were performed in the same way as described above.

For the characterization of SSR markers, the following genetic diversity indices were calculated for each locus using GenAlEx version 6.5 (Peakall and Smouse, 2012): the number of alleles (*Na*), the effective number of alleles (*Ne*), observed heterozygosity ($H_O$) and expected heterozygosity ($H_E$). Amplified products were successfully obtained using 11 primers in the multiplex PCR, and these products exhibited polymorphism (Table. S3).
